## Supplement for "Drug specificity and affinity are encoded in the probability of cryptic pocket opening in myosin motor domains"

This PDF includes:

Figures S1 to SX
Table S1 to S3

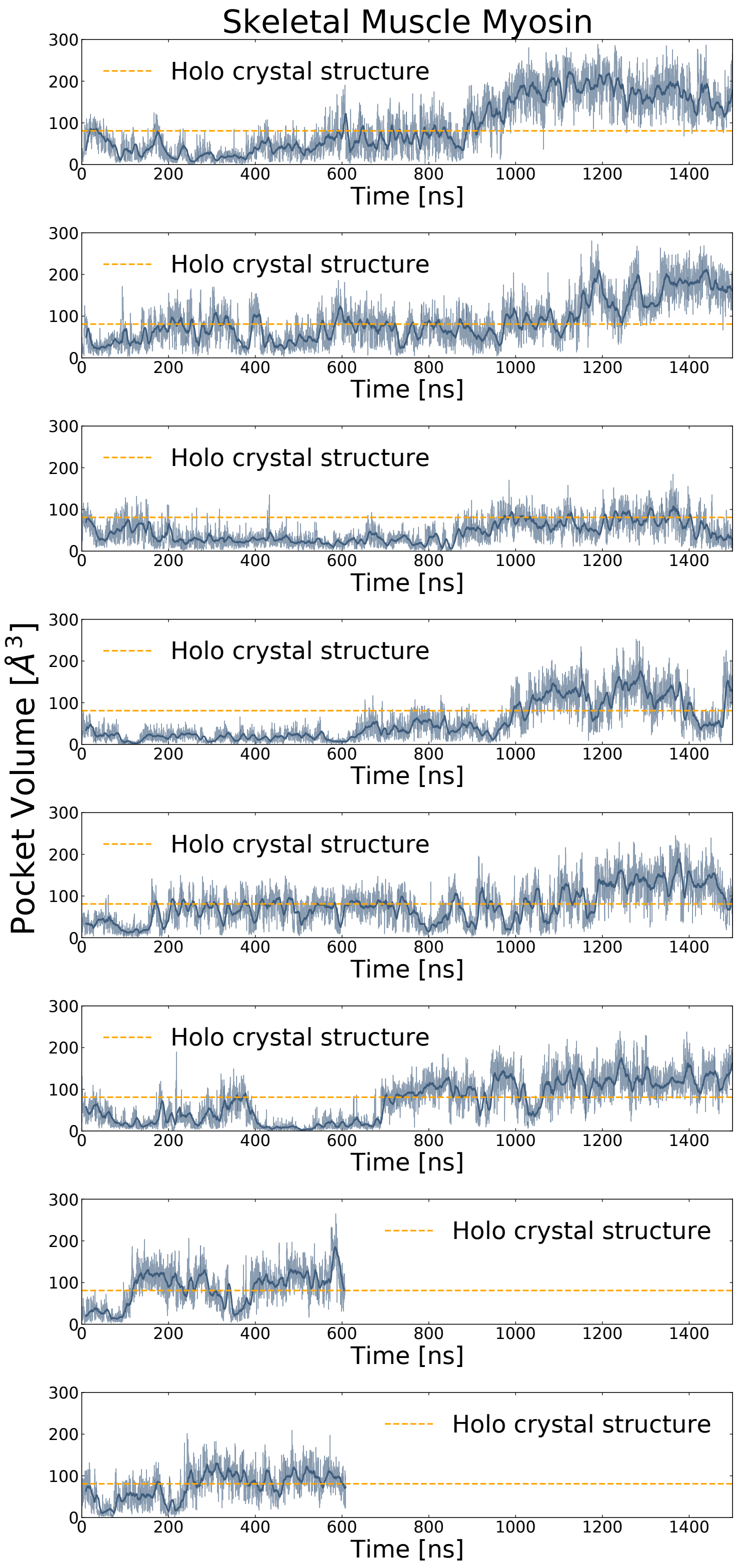


**Fig S1: Trajectory traces for long simulations of skeletal muscle myosin reveal opening in all long MD trajectories.**

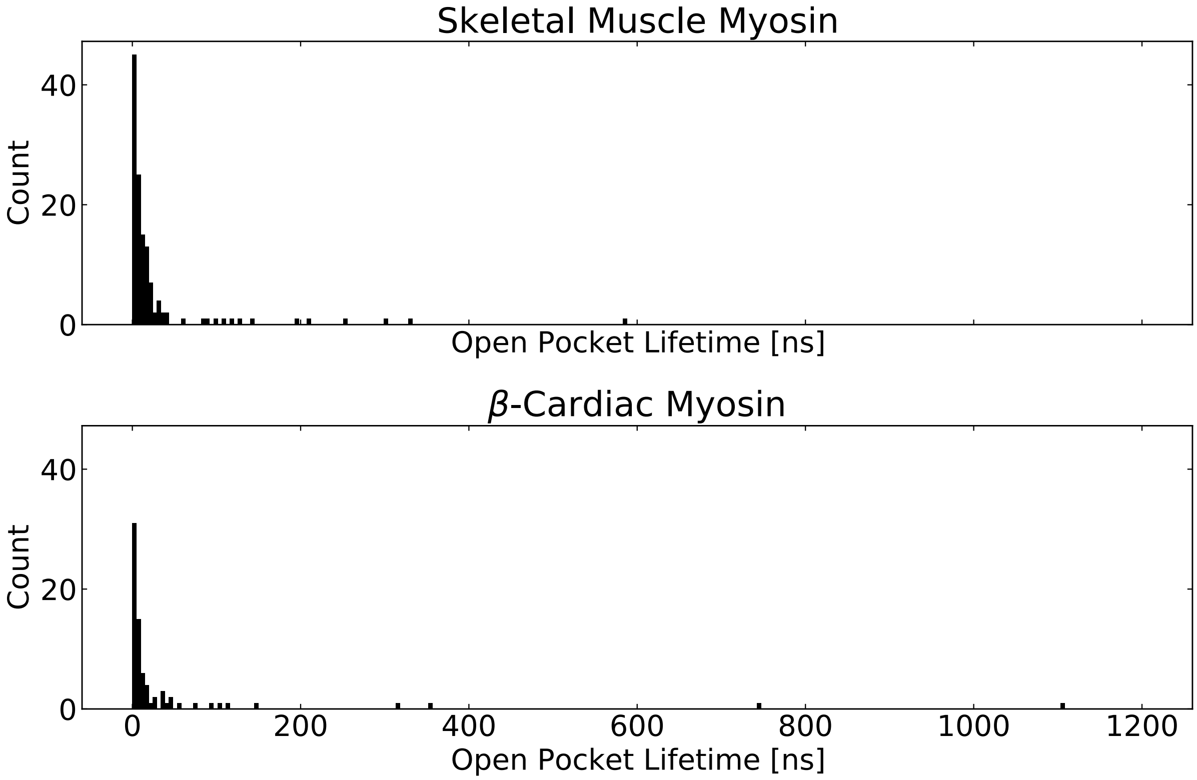


**Fig S2: The blebbistatin pocket stays open for prolonged periods (>200 ns) of simulation time in both skeletal and β-cardiac myosin**. We determined the open pocket lifetime from trajectory traces of pocket volume. Specifically, we took a 10 ns window average and determined how long the pocket volume exceeded that of the *holo* structure (PDB: 1YV3) for each opening event (i.e. trajectory time from opening to closing). For smooth muscle myosin and nonmuscle myosin 2A, a 10 ns rolling window average never exceeded the *holo* volume, though individual structures do.

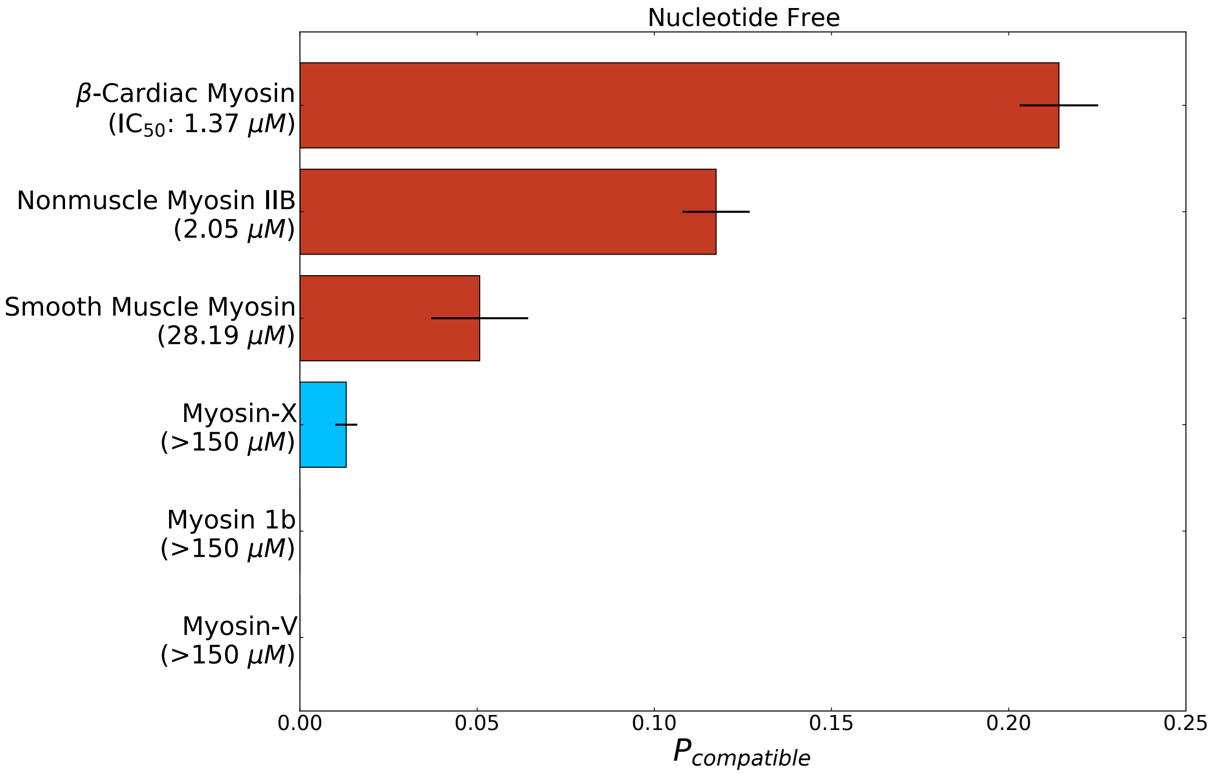


**Fig S3. The probability of adopting open pocket conformations is greater among blebbistatin-sensitive isoforms (red bars) than insensitive isoforms (blue bars)**. Structural states were considered compatible if the pocket volume at the blebbistatin binding site matched or exceeded that of a *holo* crystal structure (PDB: 1YV3). Error bars represent the standard error of the mean from 250 trials of bootstrapping where trajectories were drawn with replacement from the entire dataset.

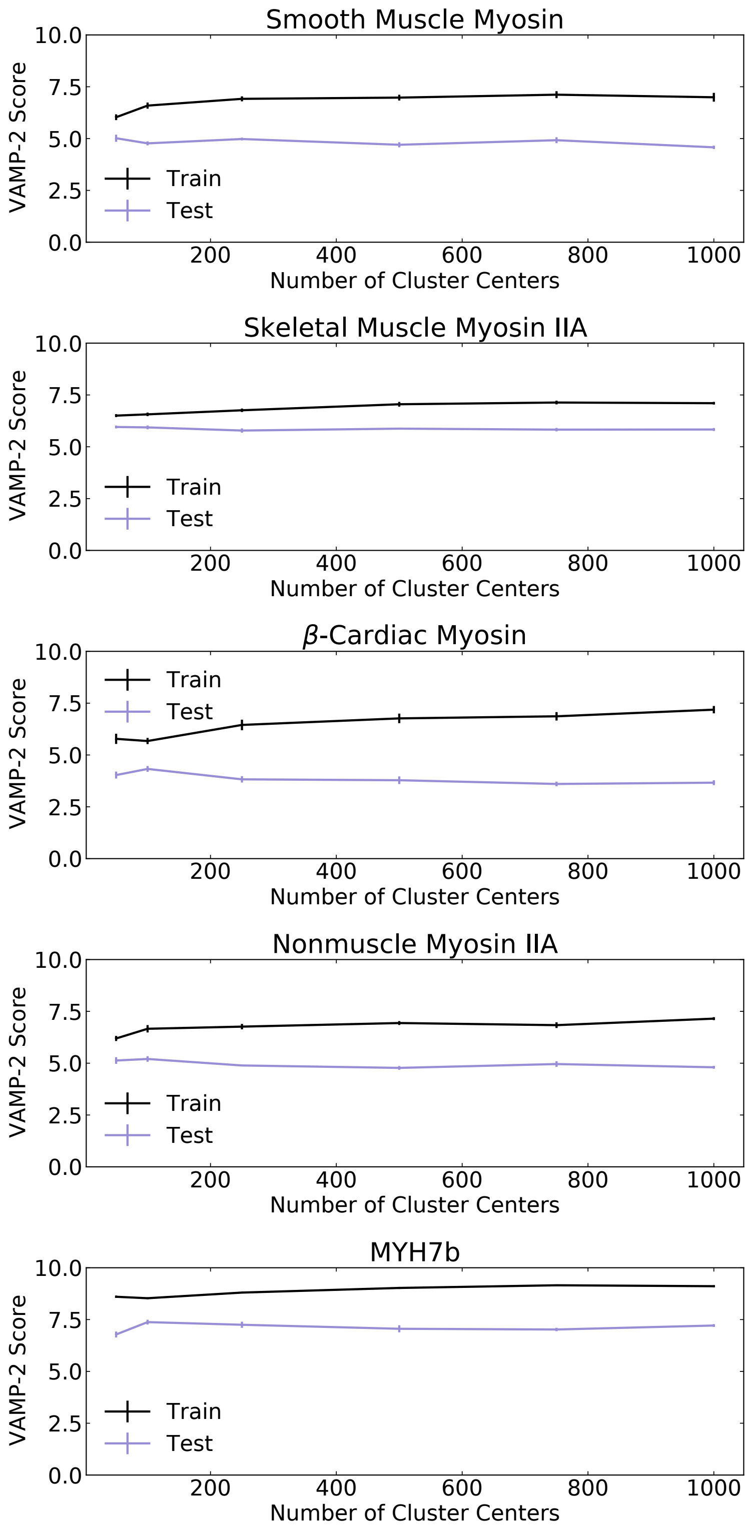


**Fig S4: VAMP-2 (Variational approach for Markov processes) scores for MSMs** constructed with varying numbers of cluster centers were computed on a validation set of trajectories to select an appropriate number of cluster centers for each MSM. Error bars show standard error of mean VAMP-2 score.

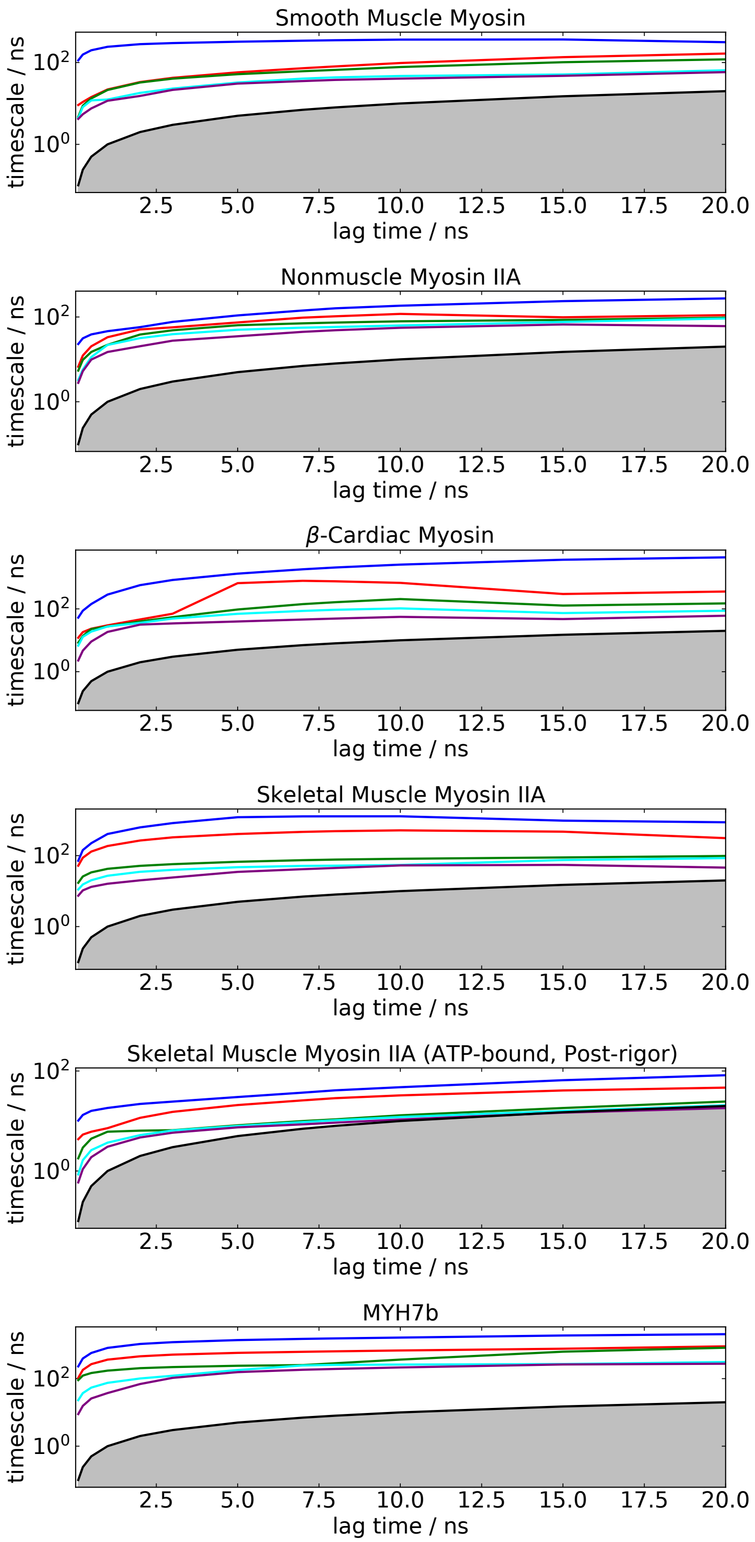


**Fig S5: Implied timescales for Markov State Models of the blebbistatin pocket** across multiple myosin isoforms show convergence on a logarithmic scale. The gray area indicates the region where timescales become equal to or smaller than the lag time and can no longer be resolved.

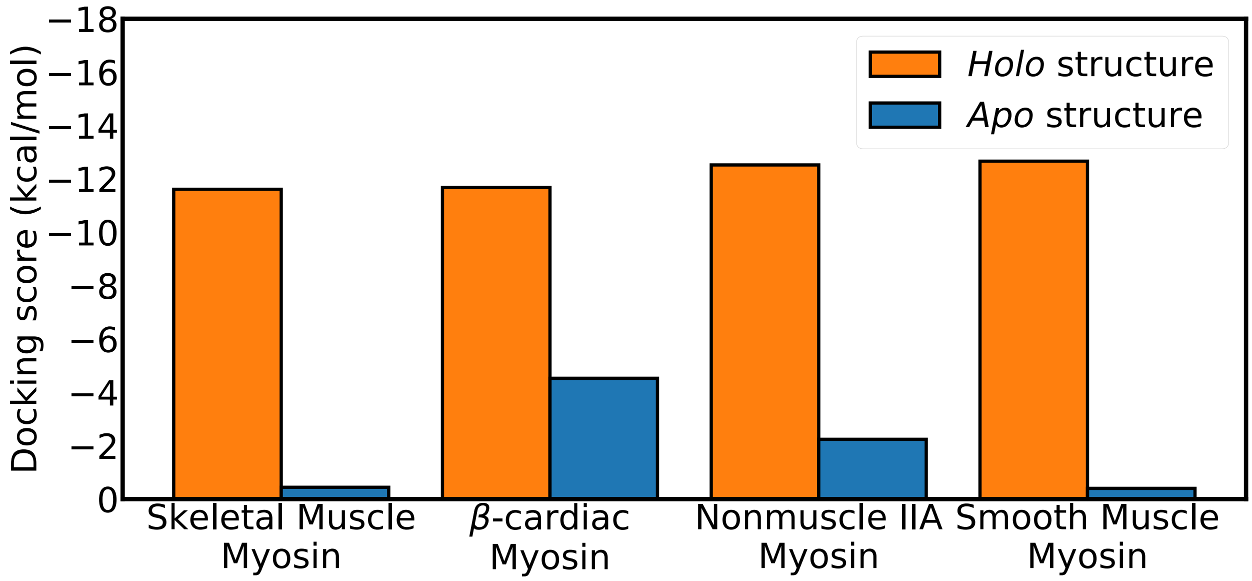


**Fig S6**: **Docking scores to homology models of *apo* and *holo* structures do not correlate with blebbistatin potency.**

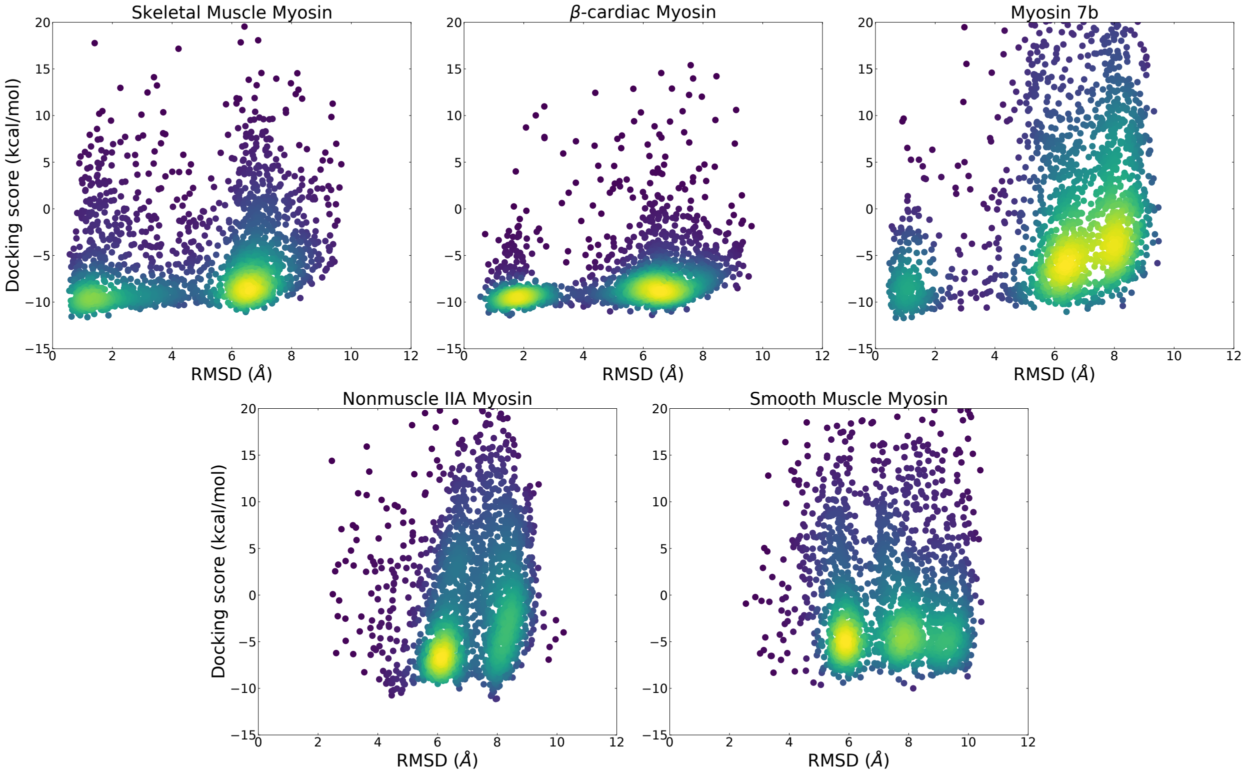


**Fig S7**: **Comparison of distribution of docking scores and ligand heavy atom root mean square deviation (RMSD)** from blebbistatin's pose in a *holo* crystal structure (PDB: 1YV3) show that skeletal and β-cardiac myosin are more likely to adopt structures where blebbistatin can be docked in its *holo* orientation and obtain a favorable docking score. Points are colored by density with bright colors indicate areas of high density.

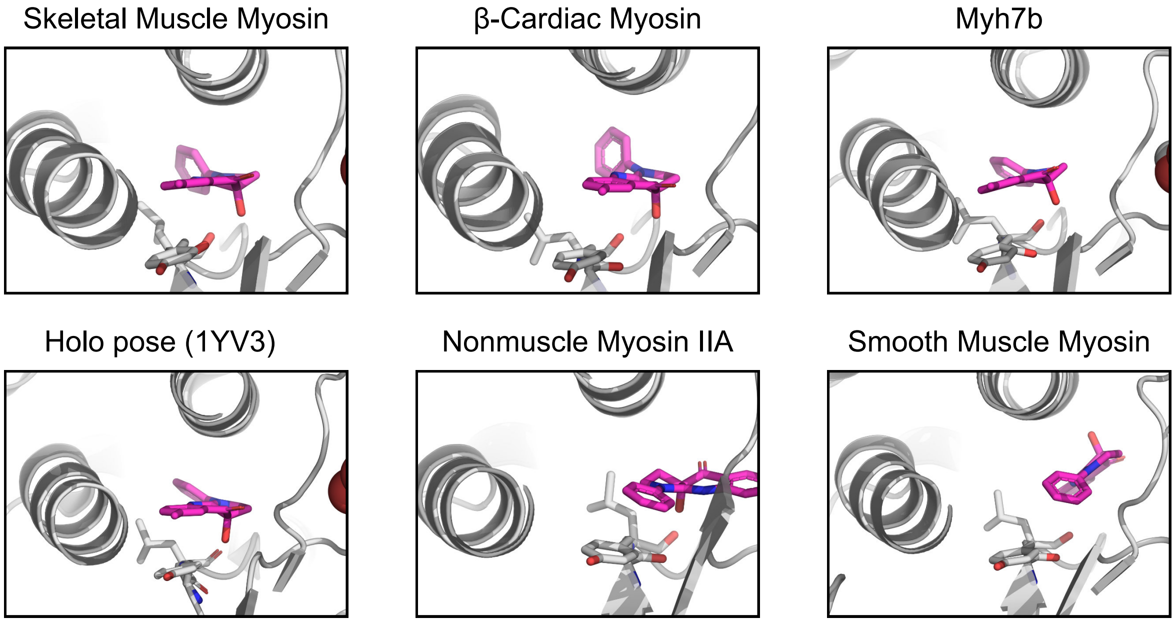


**Fig S8**: **Highest scoring poses for each of the myosin isoforms** reveals that the best pose for skeletal muscle myosin, β-cardiac myosin, and Myh7b closely matches the pose seen in *holo* crystal structures (ligand heavy atom RMSD 1.2 Å, 1.5 Å, and 0.9 Å to *holo* PDB 1YV3 for skeletal muscle myosin, β-cardiac myosin, and Myh7b respectively).

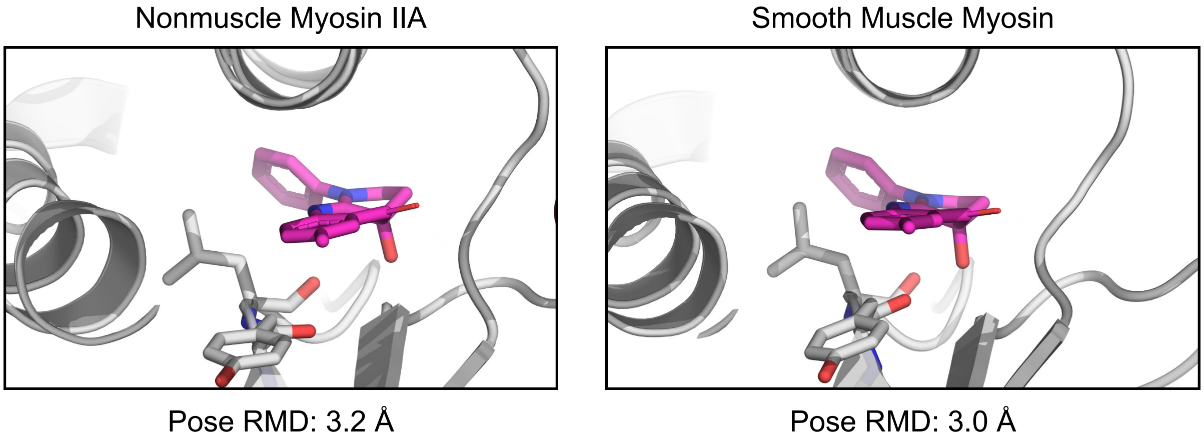


**Fig S9**: **Docking to nonmuscle myosin IIA and smooth muscle myosin produces low RMSD poses** (3.2 Å and 3.0 Å blebbistatin heavy atom RMSD from holo 1YV3 structure) with reasonably high scores (-6.3 kcal/mol in both cases).

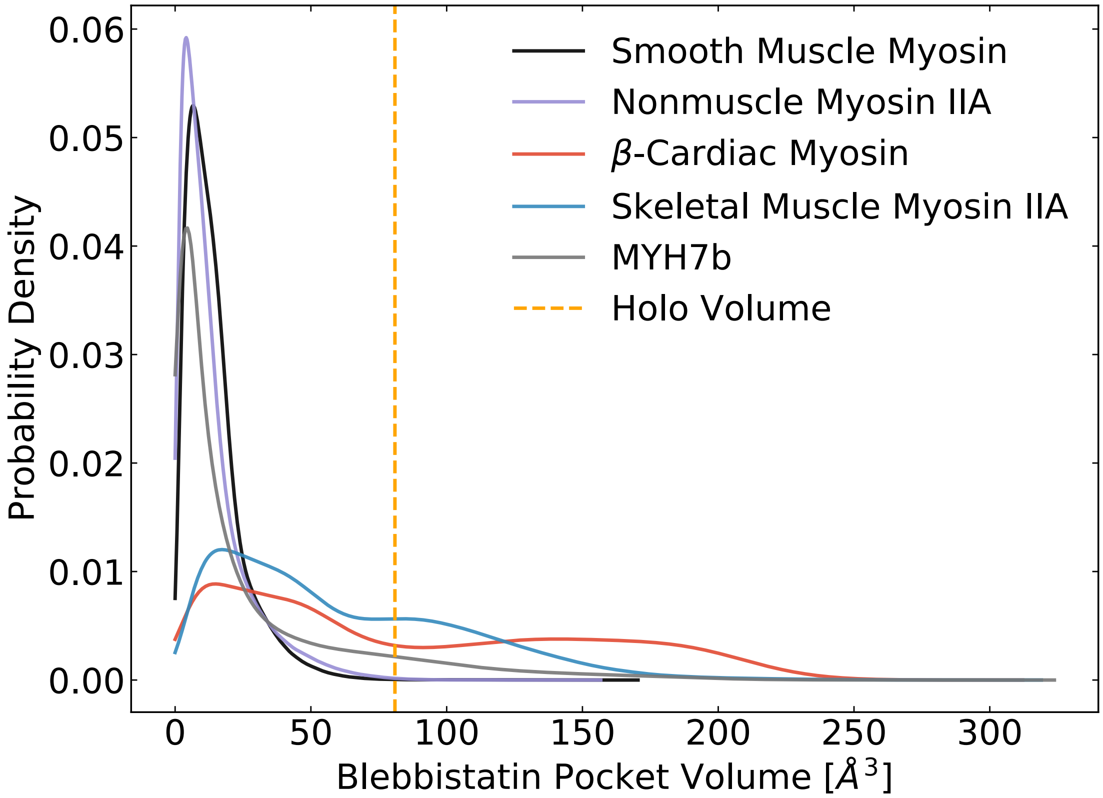


**Fig S10**: **MSM-weighted pocket volumes for myosin-II isoforms in the ADP*Pi state** reveal that blebbistatin pocket opening commonly occurs in skeletal muscle myosin, β-cardiac myosin, and Myh7b**.**

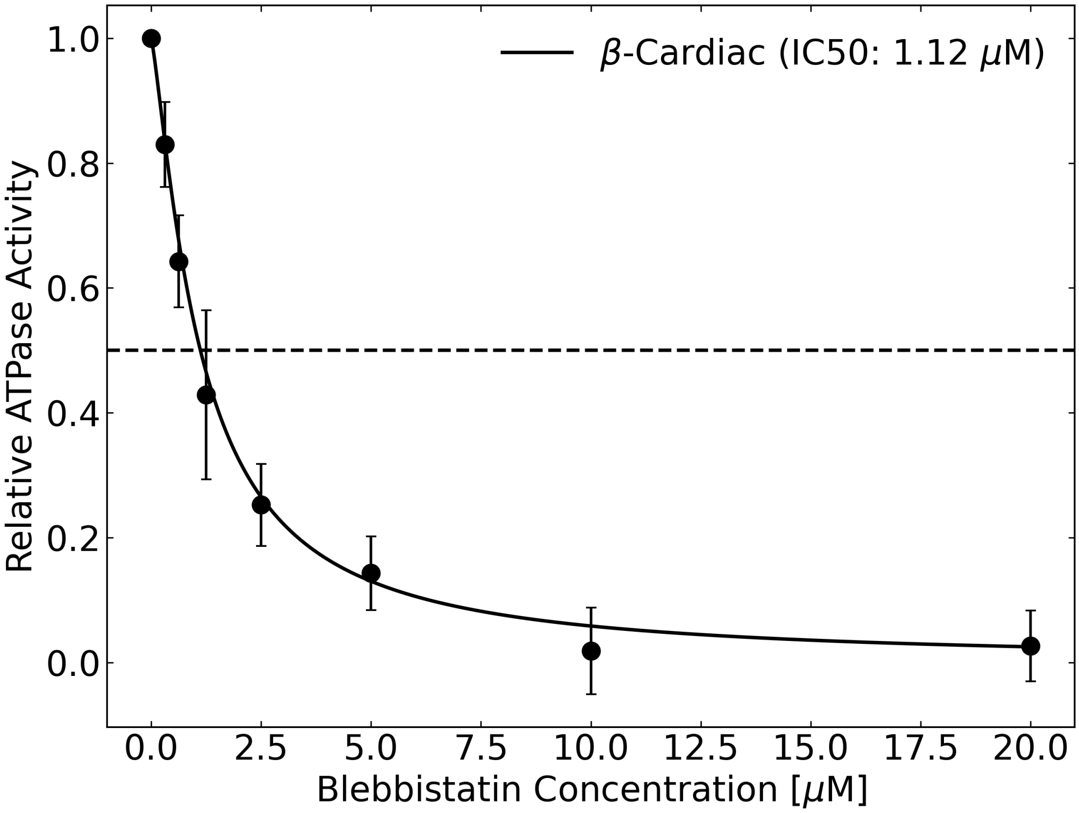


**Fig S11:** Blebbistatin inhibition of actin-activated ATPase activity was measured using an NADH-linked assay for β-cardiac myosin and an IC50 of 1.12 μM was determined. Error bars represent standard deviation across 4 trials.

### Table S1: IC50 values for different myosin isoforms

| **Isoform** | **Species** | **IC50/Ki** | **Citation** |
|---|---|---|---|
| Skeletal Muscle Myosin II | Rabbit | 0.11 | Varkuti et. al. |
| Fast skeletal | Rabbit | 0.5 | Limouze et. al. |
| Skeletal Muscle Myosin II | Rabbit | 0.28 | Radnai et. al. |
| β-cardiac | Porcine | 1.2 | Limouze et. al. |
| β-cardiac | Porcine | 1.9 | Radnai et. al. |
| Nonmuscle Myosin IIA | Human | 5.1 | Limouze et. al. |
| Nonmuscle Myosin IIA | Human | 3.58 | Zhang et. al. |
| Nonmuscle Myosin IIA | Human | 2.9 | Radnai et. al. |
| Unphosphorylated Smooth Muscle Myosin II | Chicken | 17.5 | Wang et. al. |
| Unphosphorylated Smooth Muscle Myosin II | Bovine | 10.1 | Wang et. al. |
| Phosphorylated Smooth Muscle Myosin II | Chicken | 23.5 | Wang et. al. |
| Smooth Muscle Myosin | Chicken | 6.47 | Zhang et. al. |
| Smooth Muscle Myosin 2A and 2B | Chicken | 3 | Eddinger et. al. |
| Smooth Muscle Myosin | Turkey | 79.6 | Limouze et. al. |
| Smooth Muscle Myosin II | Chicken | 3.2 | Radnai et. al. |

**Table S2: Summary of simulations performed for this study and MSM hyperparameters**

| System | Structural State | Number of Cluster Centers | Lag Time (ns) | Total Simulation Time (μs) | Median Trajectory Length (ns) | Maximum Trajectory Length (ns) |
|---|---|---|---|---|---|---|
| Skeletal Muscle Myosin | ADP*Pi | 50 | 5 | 89.0 | 21.2 | 1500 |
| β-Cardiac Muscle Myosin | ADP*Pi | 100 | 5 | 90.1 | 21.0 | 1500 |
| Nonmuscle Myosin IIA | ADP*Pi | 100 | 8 | 86.7 | 20.0 | 1500 |
| Smooth Muscle Myosin | ADP*Pi | 50 | 5 | 87.0 | 20.0 | 1500 |
| Skeletal Muscle Myosin | ATP | 100 | 5 | 2.7 | 910.0 | 925 |
| Myosin 7b | ADP*Pi | 100 | 5 | 100.8 | 375.0 | 1500 |

**Table S3: Percent identity in motor domain sequence between myosin-II isoforms in this study**

| | MYH11 | MYH9 | MYH7b | MYH7 | MYH2 |
|---|---|---|---|---|---|
| MYH11 | 100 | 84.58 | 52.03 | 52.45 | 51.56 |
| MYH9 | 84.58 | 100 | 51.01 | 52.23 | 50.67 |
| MYH7b | 52.03 | 51.01 | 100 | 69.65 | 66.24 |
| MYH7 | 52.45 | 52.23 | 69.65 | 100 | 80.77 |
| MYH2 | 51.56 | 50.67 | 66.24 | 80.77 | 100 |
